# Rapamycin Reduces Mineral Density and Promotes Beneficial Vascular Remodeling in a Murine Model of Severe Medial Arterial Calcification

**DOI:** 10.1101/2024.08.01.606196

**Authors:** Parya Behzadi, Andrew A. Wendling, Rolando A. Cuevas, Alex Crane, Claire C. Chu, William J Moorhead, Ryan Wong, Mark Brown, Joshua Tamakloe, Swathi Suresh, Payam Salehi, Iris Z. Jaffe, Allison L. Kuipers, Lyudmila Lukashova, Konstantinos Verdelis, Cynthia St. Hilaire

**Affiliations:** Department of Medicine, Division of Cardiology, and the Pittsburgh Heart, Lung, Blood and Vascular Medicine Institute, University of Pittsburgh, Pittsburgh, Pennsylvania, USA; CardioVascular Center, Vascular Surgery, Tufts Medical Center, 800 Washington Street, Boston, MA, 02111-1800, USA; Molecular Cardiology Research Institute, Tufts Medical Center, 800 Washington Street, Boston, MA, 02111-1800, USA; Departments of Medicine and Epidemiology & Biostatistics, College of Human Medicine, Michigan State University, Grand Rapids, MI, USA; Departments of Endodontics and Oral Biology, School of Dental Medicine, University of Pittsburgh, Pittsburgh, PA, USA; Department of Bioengineering, University of Pittsburgh, Pittsburgh, Pennsylvania, USA; Department of Cardiothoracic Surgery, University of Pittsburgh, Pittsburgh, Pennsylvania, USA

**Keywords:** Medial Arterial Calcification, Rapamycin, Matrix GLA protein, Collagen, Elastin

## Abstract

Peripheral artery disease (PAD) is the narrowing of the arteries that carry blood to the lower extremities. PAD has been traditionally associated with atherosclerosis. However, recent studies have found that thrombotic events triggered by medial arterial calcification (MAC) is the primary cause of chronic limb ischemia below the knee. MAC is localized around the elastic fibers surrounding smooth muscle cells (SMCs) in arteries. Matrix GLA protein (MGP) binds circulating calcium and prevents hydroxyapatite mineral deposition, while also modulating pro-osteogenic signaling by attenuating BMP-2-mediated activation of *Runx2* gene expression. *Mgp*^-/-^ mice develop severe MAC and die around 8 weeks after birth due to aortic rupture or heart failure. We previously discovered a rare genetic disease Arterial Calcification due to Deficiency of CD73 (ACDC), in which patients present with extensive MAC in their lower extremity arteries. Using a patient-specific induced pluripotent stem cell model, we found that rapamycin inhibited calcification. Here we investigated whether rapamycin could reduce MAC in vivo using the *Mgp*^-/-^ murine model. *Mgp*^*+/+*^ and *Mgp*^-/-^ mice received 5mg/kg rapamycin or vehicle. Calcification content was assessed via microCT, and vascular morphology and extracellular matrix content were assessed histologically. Immunostaining and western blot analysis were used to examine SMC phenotype and extracellular matrix content. Rapamycin prolonged *Mgp*^-/-^ mice lifespan, decreased mineral density in the arteries, maintained SMC contractile phenotype, and improved vessel structure, however, calcification volume was unchanged. *Mgp*^-/-^ mice with SMC-specific deletion of Raptor or Rictor did not recapitulate treatment with rapamycin. These findings suggest rapamycin promotes beneficial vascular remodeling in vessels with MAC.

**NEWS AND NOTEWORTHY:** Peripheral artery disease (PAD) is associated with medial arterial calcification (MAC), which involves calcification of arterial elastic fibers and smooth muscle cells (SMCs). Matrix GLA protein (MGP) inhibits vascular calcification, and *Mgp*^*-/-*^ mice develop severe MAC. Using this model, we found rapamycin prolonged lifespan, reduced arterial mineral density, maintained SMC contractile phenotype, and improved vessel structure, though calcification volume remained unchanged. Findings highlight rapamycin’s potential for vascular remodeling in MAC.

## INTRODUCTION

Peripheral artery disease (PAD) is the narrowing of blood vessels in the lower extremities due to inward remodeling or thrombotic occlusion, both of which cause chronic limb ischemia and often result in amputation of the lower leg (1). Traditionally atherosclerosis has been the assumed cause of PAD. However, recent studies now show that medial arterial calcification (MAC) promotes inward remodeling, and stiffness related to medial dysplasia induces thrombosis (2-4). MAC is characterized by the progressive buildup of calcium and phosphate along the elastic fibers within the arterial walls and is typically not associated with lipid deposition or fibrous cap formation (5). These structural features distinguish PAD from atherosclerosis and indicate that MAC is a distinct pathology driving adverse outcomes in PAD. Currently, there are no specific medical therapies that target the distinct pathogenesis of PAD or MAC. Finding therapies that can prevent, stop, or even reverse MAC in PAD could greatly enhance the current standard of care.

Vascular calcification stems from the nucleation of calcium and phosphate into hydroxyapatite crystals and the release of pro-mineralizing matrix vesicles from osteogenic SMCs that accumulate along the elastic lamina (6). Phosphate is a byproduct of the breakdown of extracellular adenosine triphosphate (ATP) to adenosine, generating inorganic phosphate at several points (7, 8). There are several genetic diseases that present with MAC, which harbor inactivating mutations in the genes related to extracellular ATP metabolism (5). Patients with Arterial Calcification due to Deficiency of CD73 (ACDC, OMIM # 211800) harbor inactivating mutations in the *NT5E* gene, which encodes for the CD73 enzyme that primarily metabolizes extracellular AMP to adenosine and inorganic phosphate. Key signatures of this disease are calcification nodules in the small joint capsules and extensive MAC in the lower-extremity arteries that initiate along the elastic fibers (4, 9, 10). We previously found that a lack of CD73-mediated adenosine enhanced the expression and activity of tissue-nonspecific alkaline phosphatase (TNAP), a key enzyme that promotes calcification (11, 12), as well as altered elastin deposition and SMC contractility in response to TGF-β (13).

Matrix GLA protein (Mgp) is a vitamin-K2-dependent protein with an unusual gamma-carboxylation of five glutamate residues that enhance its affinity for calcium while preventing it from mineralizing (14). Mgp inhibits calcification propagation by binding hydroxyapatite crystals and stimulating their uptake by local phagocytosing macrophages (15). Mgp also modulates osteogenic signaling by binding to BMP-2, preventing it from activating receptors and upregulating Runx2, the key regulator of osteoblast differentiation and maturation (16, 17). Mice that lack *Mgp* develop to term but die prematurely within approximately two months of age due to extensive MAC, which leads to aortic rupture or heart failure (18). Aortas of *Mgp*-deficient mice exhibit increased collagen accumulation and elastic fiber fragmentation within the medial layer of the arteries, where calcification is also localized. This phenocopies MAC observed in human patients with PAD MAC and ACDC patients (4).

Rapamycin binds the FK506-binding protein (FKBP12), converting it to a potent allosteric inhibitor of the mammalian target of rapamycin (mTOR) complex. mTOR is broadly expressed throughout the body, which controls many key processes such as energy balance, autophagy, and proliferation (19-22). Rapamycin has been found to protect against the calcification of vascular cells in in vitro and ex vivo models, as well as in in vivo models of chronic kidney disease-induced MAC (23-25). We previously found rapamycin prevented calcification in in vitro and in in vivo disease models of ACDC patient-specific induced pluripotent stem cells (iPSCs) (11).

While CD73-deficient humans develop extensive MAC and broken elastin fibers in their lower extremity large vessels, CD73-deficient mice do not phenocopy humans and do not exhibit calcification in their vasculature (26, 27). As the MAC observed in the aorta *Mgp*-deficient mice mirrors that seen in the affected vessels of ACDC patients, we sought to investigate whether rapamycin prevented calcification in this genetic mouse model of MAC.

## MATERIALS AND METHODS

### Availability of Materials

We abide by the NIH Grants Policy on Sharing of Unique Research Resources, including the NIH Policy on Sharing of Model Organisms for Biomedical Research (2004), NIH Grants Policy Statement (2003), and Sharing of Biomedical Research Resources: Principles and Guidelines for Recipients of NIH Grants and Contracts (1999), and the Bayh-Dole Act and the Technology Transfer Commercialization Act of 2000. Materials generated in our laboratory are made available for non-commercial research per established University of Pittsburgh Office of Research IRB and MTA protocols.

### Human tissue collection

De-identified human tissues were obtained, with informed consent from subjects, who were enrolled in studies approved by the institutional review board (IRB) of the University of Pittsburgh or Tufts Medical Center, per the Declaration of Helsinki. Personnel involved with specimen handling underwent all required institutional training. Human coronary artery and tibial artery tissues were excised and washed with a sterile rinsing solution (sterile PBS supplemented with 2.5 μg/mL of fungicide (Gibco, 15290026), 0.05 mg/mL of gentamicin (Gibco, 15710064), and 5 μg/mL of plasmocin (InvivoGen, ant-mpt-1). Tissues were then fixed in 4% paraformaldehyde in PBS for 2 hours, placed in PBS, embedded in paraffin blocks for pathology, and cut to a thickness of 10 μm.

### Animals and cell line generation

Animal use was approved by the Institutional Animal Care and Use Committee at the University of Pittsburgh. Mice were under the veterinary care of the University of Pittsburgh Division of Laboratory Animal Resources, which adheres to the NIH policy on the Animal Welfare Act and all other applicable laws. Facilities are under the full-time supervision of veterinarians and are AAALAC-accredited. Our protocols follow the AVMA Guidelines on Euthanasia. *Mgp* heterozygous mutant mice from a C57BL/6J background (strain# 023811 Jackson Laboratory, Bar Harbor, ME) were bred to produce wild-type (+/+) and knockout (-/-) littermate controls. Smooth muscle lineage-specific *Raptor* (*Mgp*^*-/-*^; *Raptor*^*SMC-/-*^) and *Rictor* (*Mgp*^*-/-*^; *Rictor*^*SMC-/-*^) knockout mice were produced by breeding *Mgp*^*+/-*^ with Myh11-Cre-eGFP mice (strain# 007742 Jackson Laboratory, Bar Harbor, ME)(28), with Raptor-LoxP (strain# 013188 Jackson Laboratory, Bar Harbor, ME)(29), or *Rictor*-LoxP (strain# 020649 Jackson Laboratory, Bar Harbor, ME)(30). Genotypes were determined from tail snips that were incubated in DirectPCR Lysis Reagent (Viagen Biotech) according to manufacturer instructions. The lysate was amplified using OneTaq MasterMix (New England Biolabs) according to manufacturer instructions with primers suggested by Jackson Laboratories.

### Cell culture

SMC lines were generated from three-to five-week-old male and female *Mgp*^*+/+*^ and *Mgp*^-/-^ mice according to previously described protocols (13). *Mgp*^*+/+*^ and *Mgp*^-/-^ aortic SMC lines were cultured in Dulbecco’s modified Eagle’s medium (DMEM; Gibco, Waltham, MA) supplemented with 20% FBS (FBS; R&D Systems, Minneapolis, MN) and 100 U/mL penicillin-streptomycin (P/S; Gibco, Waltham, MA). Growth media was changed every three days, and cells were split 1:2 when confluent. Post-expansion, 25,000–50,000 cells/cm^2^ were plated in DMEM supplemented with 10% FBS and 100 U/mL penicillin-streptomycin, and grown to confluence. Before all experiments/treatments, cells were serum-starved in DMEM with 0.5% FBS for 48 h. Incubation under reduced serum conditions has been proven to be a beneficial technique for studying smooth muscle cell remodeling and contractility in vitro (31, 32). Cells were treated with rapamycin (Novus Biologicals) at concentrations of 200 nM or the same volume of DMSO as vehicle control for 14h. For autophagy flux assessment, cells were exposed to either bafilomycin A1 100 nM (Sigma-Aldrich) or rapamycin 200 nM for a duration of 24h following treatment intervention. All chemicals were dissolved in DMSO and administered in equal volume as vehicle control.

### Murine Tissue Extraction

Mice were sacrificed by asphyxiation with carbon dioxide, followed by perforation of the diaphragm. Tissue extraction began with dissection through the abdominal wall and then perfusion of the heart with 50 mL of DPBS supplemented with 1:100 Amphotericin B (Gibco,15290026) and 1:200 Gentamicin (Gibco, 15710064). The aortas were maintained in this solution while the adventitia was removed in a petri dish. Tissues were then fixed in 4% paraformaldehyde in PBS for 2 hours and then placed in PBS. Tissues were brought to microCT and then embedded in paraffin and cut to a thickness of 10 μm.

### In Vivo Rapamycin Injections

Male and female *Mgp*^+/+^ and *Mgp*^-/-^ mice received intraperitoneal injections (1x or 3x per week) of either vehicle or 5 mg/kg rapamycin dissolved in 0.3 mg/ml peanut oil beginning at 10 days of age. This process continued until the natural death or a collection timepoint.

### Micro-Computed Tomography

Five-10mm in length segments of mouse arteries sealed in parafilm and stabilized inside a holder were imaged by a Scanco µCT 50 (Scanco Medical, Brüttisellen, Switzerland) microcomputed tomography system with a 3.4 µm voxel resolution, with a 55 KVp beam energy, and 145 µA (1500 ms exposure per view). The Scanco µCT software (HP, DECwindows Motif 1.6) was used for 3D volume reconstruction, and the Scanco morphometry and densitometry software was used for the processing of images. Measurements included the total volume and density of mineral deposits after segmentation from the background with a 200 mg/cc threshold.

### Western Blot Analysis

Cells were lysed in 1% CHAPS hydrate, 150 mmol/L sodium chloride, and 25 mmol/L HEPES buffer supplemented with 1× protease and phosphatase inhibitor (Sigma-Aldrich). Cells were scraped into microcentrifuge tubes, vortexed for 5 minutes, frozen/thawed for 5 to 8 cycles, then centrifuged at 12□000×g for 10 minutes at 4°C. Supernatant protein concentration was determined using Pierce BCA protein assay kit (Thermo Fisher, Waltham, MA). Ten micrograms of protein were used to prepare lysate with 1 × Pierce LDS sample buffer nonreducing (Thermo Fisher, Waltham, MA) and 1 × NuPAGE sample reducing agent (Novex, Waltham, MA). Lysates were denatured at 95°C for 15 min and then electrophoresed on 4%–20% TGX stain-free polyacrylamide gel (Bio-Rad, Hercules, CA) in 1 × Tris/Glycine/SDS buffer (Bio-Rad, Hercules, CA) at 120 V for 50 min. Protein was transferred onto a 0.2 µm nitrocellulose membrane in prepared 1 × Towbin buffer with 100% ethanol (EtOH) at 1 A and 25 V for 30 min using the Trans-Blot Turbo Transfer System (Bio-Rad, Hercules, CA). Membranes were blocked in 1:1 Odyssey blocking buffer (Li-COR, Lincoln, NE) and PBS for 1 h at room temperature, followed by primary antibodies incubation of LC3 (Abcam, ab192890, RRID: AB_2827794) and α-tubulin (926-42213, LI-COR, RRID: AB_2637092) in 1:1 Intercept blocking buffer (LI-COR, 927-70001) plus 0.1% Tween 20 (PBS-T) at 4°C overnight. Membranes were washed in PBS-T three times for 5 min, then incubated in secondary antibodies, anti-rabbit(LI-COR, 926-32211, RRID: AB_621843) or anti-mouse antibody (LI-COR, 926-32210, RRID: AB_621842), at room temperature for 1 h. Membranes belonging to the same experimental set were imaged simultaneously on an Odyssey CLx (LI-COR, Lincoln, NE), and band intensity quantification was performed with Image Studio (Version 5.2, LI-COR, Lincoln, NE) software. Individual bands were normalized to α-tubulin, and each treatment group’s fold change was compared with each gel’s vehicle control lanes. For sequential antibody incubations, membranes were stripped in 1 × NewBlot Nitro Western Blot Stripping Buffer (LI-COR, Lincoln, NE) for 10 min, followed by three washes in PBS.

### Verhoeff Van Gieson, Von Kossa, Masson’s Trichrome and Picrosirius Red Staining

Mice aortic tissues were removed from mice through the abdominal wall under anesthesia conditions. Then aortic tissue was fixed with 4% paraformaldehyde and embedded in paraffin. The embedded specimens were transversely sectioned at 10 µm on a microtome cryostat (Microm HM 325). Slides with the adhered paraffin aortic sections were warmed to 65°C for 1 h and then deparaffinized through xylene, rehydrated with serial incubation in graded alcohol baths, and stained with Verhoeff–van Gieson for elastic fiber visualization (Polysciences, 25089-1), Von Kossa for calcification (Polysciences, 24633-1), Masson’s for collagen fibers (Polysciences, 25088-1) and Picrosirius Red staining (Polysciences, 24901-500) for collagen quantification according to manufacturer’s instructions. Verhoeff Van Gieson, Von Kossa, Masson’s Trichrome images were captured using Fritz PreciPoint scanner with 20x Nikon objective. Collagen staining was assessed by measuring the intensity of collagen in the vessel wall normalized to scale bar. Picrosirius Red staining images captured collagen with polarized transmitted light microscopy (Nikon Eclipse NI-E) stand with a plan Apo 20x Nikon objective and a Teledyne Prime BSI Express camera with an exposure time of 100msec. The percentage of collagen in the medial layer was measured using Nikon Elements 6.02.03. The percentage of medial thickness of the medial layer was measured using ImageJ software (Java 8), where the medial thickness was determined by measuring the distance between the inner and outer elastic lamina at four sites per vessel. The percentage of calcification in the medial layer was measured using ImageJ software (Java 8) where the area of calcification was normalized to total medial area. The tortuosity of elastic fibers was measured by selecting three representative areas with the whole vessel. ImageJ was used to trace and measure the length of five elastic fibers and shown as a ratio over the vessel length. The number of breakages in the elastic lamina was assessed by counting the breaks throughout the entire aorta. All length measurements were normalized to the scale bar units.

### Immunofluorescence

Slides with the adhered paraffin aortic sections were warmed to 65°C for 1 h and then deparaffinized through xylene, rehydrated with serial incubation in graded alcohol baths. Slides were boiled in citric acid-based antigen retrieval solution (H-3300, Vector Labs) for 20 minutes. Slides were cooled in the unmasking solution for 1 hour, washed in 1x PBS and placed in blocking buffer (500 mL PBS 0.3g Fish Skin Gelatin and 10% horse serum) for 1 hour. Tissues were incubated overnight at 4°C with antibodies for α-SMA (Abcam, ab5694, RRID: AB_2223021, 1:50), Ki67 (Abcam, ab15580, RRID: AB_443209, 1:250), LC3 (Abcam, ab192890, RRID: AB_2827794, 1:50), Mgp (Proteintech, 10734-I-AP, RRID: AB_2297660, 1:25), Myh11 (Invitrogen, MA5-4284, RRID: AB_2911986, 1:50), Runx2 (ABclonal, A2851, RRID: AB_2764676 1:100). Cells were washed 3× for 5 minutes each with PBS, 0.1% TWEEN 20, then once with PBS, and then incubated with secondary antibodies (Invitrogen, A-11012, A-11008) diluted in a blocking solution for 1 hour. Cells were washed 3× for 5 minutes each with PBS, 0.1% TWEEN 20, then once with PBS, then mounted with Fluoroshield Mounting Medium with DAPI (Ab104139, Abcam). All signals were optimized to a rabbit polyclonal IgG (Vector, I-1000, RRID: AB_2336355, 1:50) at the same concentration of the specific antibodies. Slides were imaged within 24 hours of mounting. Cells were imaged within 24 hours of mounting. Images were obtained using a Nikon NIE-I microscope at 20x objective. The measured intensity of a given stain was normalized to DAPI intensity of that same image. In ImageJ, each image was split into blue, red, and green channels, and the pixel intensity of the red and blue channels was measured (33). Representative images are selected based on quality, and if the assay includes quantification, the image that shows the average of the quantification across multiple images.

### Blinding Procedures

Professionally trained technicians handled animal welfare, including weaning and mouse line expansion, diets, sacrifice, and surgery. For experiments, blinding procedures were used, and individuals conducting and analyzing experiments did not know the treatment groups until after the analysis was performed.

### Sample Size and Statistics

Statistical analysis was performed with GraphPad Prism 9.2 (GraphPad Software, Inc) and data shown are mean ± standard deviation (SD). All experiments used at least n = 4 biological replicates and were run in technical duplicates, exact n per group specified in figure legends. Mantel-Cox test and Gehan-Breslow-wilcoxon test were used for Kaplan-Meier analysis. Statistical comparisons between the two groups were performed by either nonparametric Mann-Whitney U test or Kruskal-Wallis test when n ≤ 6 ≤ or parametric Two-way ANOVA when n > > 6. Statistical analysis used, exact n values, biological replicates, and P values are stated within each figure legend. A p-value equal to or less than 0.05 will be considered statistically significant.

## RESULTS

### MAC in MGP-deficient mice phenocopies human MAC

The MAC is localized in the medial layer of arteries along the elastic fibers. Von Kossa staining for calcification illustrates that *Mgp*^-/-^ mouse aortas have extensive mineralization in the medial layer that phenocopies the mineralization seen in MAC of the human tibial arteries, and is distinct from neointimal calcification found in the necrotic core of coronary artery with atherosclerotic plaque (Figure 1A). Mgp is a potent inhibitor of soft tissue calcification. *Mgp*^-/-^ mice do not produce functional Mgp protein (Figure 1B, 1C) and, as a result, develop extensive MAC pathology in large vessels (18).

**Figure 1:**
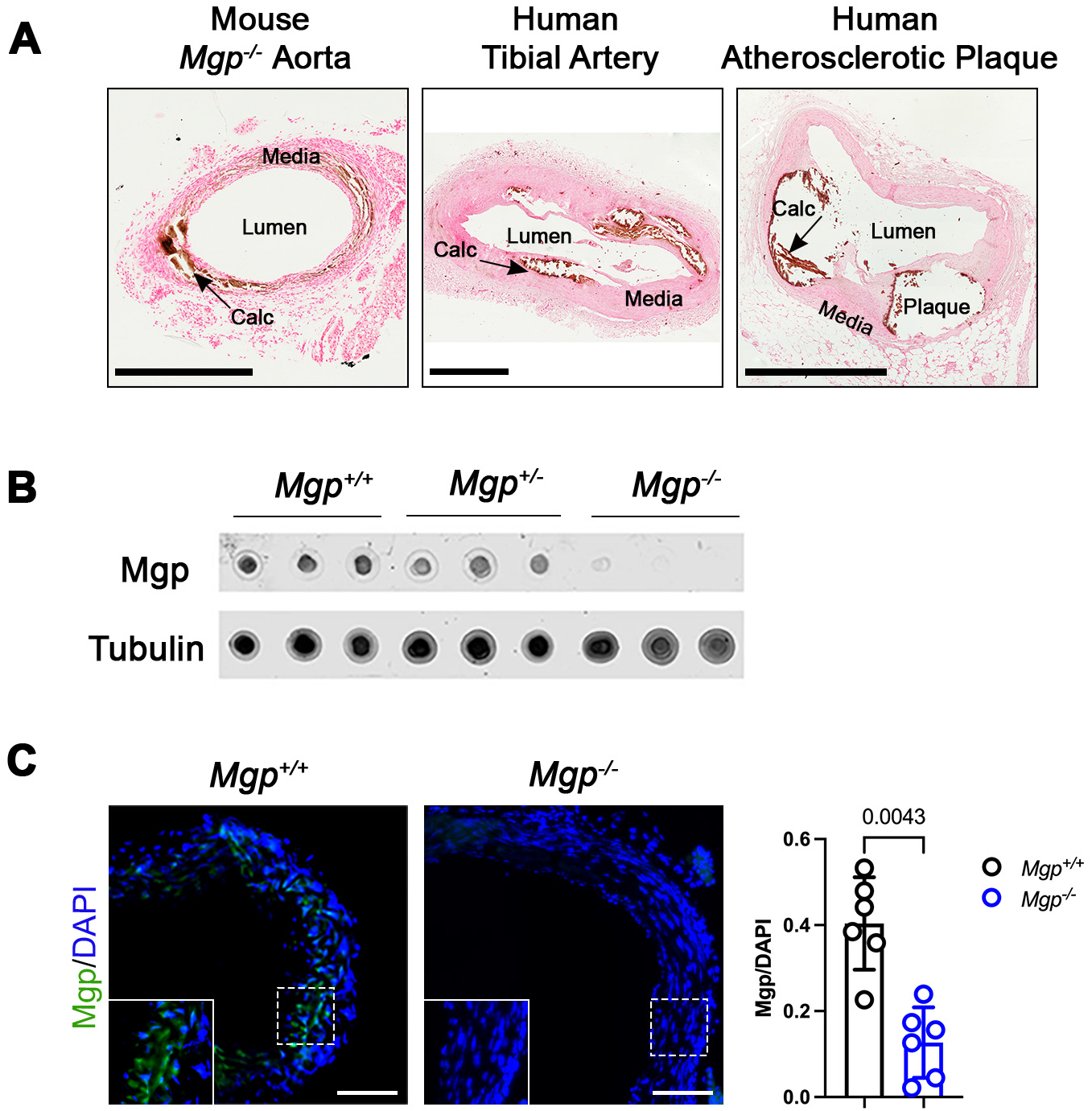
Histological comparison of Matrix Gla Protein (Mgp) mice arteries, medial and intimal calcification in human arteries. **A**. Histological sections of *Mgp*^*-/-*^ mouse aorta, human tibial plaque, and human coronary artery with atherosclerotic plaque, stained for calcification (Cal) using Von Kossa Staining. Scale Bars: human tissue 1 mm, mouse tissue 0.5 mm. **B**. Dot Blot Assay to identify Mgp protein in *Mgp*^*+/+*^, *Mgp*^*+/-*^, and *Mgp*^*-/-*^ mice. Results are representative of n=3 per group. **C**. Representative immunofluorescence images of Mgp protein in aortas from *Mgp*^*+/+*^, *Mgp*^*+/-*^ and *Mgp*^*-/-*^ mice. Scale Bars: 100 μm. n=6 *Mgp*^*+/+*^ vehicle, n=6 *Mgp*^*-/-*^ vehicle. P values were calculated using Mann-Whitney U test. Data are shown as means ± SD.

### Rapamycin prolongs lifespan and reduces arterial mineral density in *Mgp*^*-/-*^ mice

Humans with homozygous inactivating mutations in *NT5E* develop extensive MAC in their lower extremity arteries (9). While the *Nt5e*-knockout mice model shows altered renal function, diminished control of the glomerular arteriolar tone, elevated atherogenesis, elevated thrombotic occlusion, and elevated hypoxia-induced vascular leakage (26), these mice do not mimic the human vascular calcification phenotype (26, 27, 34). Jin et al. circumvented this issue using a patient-specific in vitro and in vivo disease modeling system using iPSC technology. They demonstrated that patient-specific ACDC-iPSCs developed extensive calcification in the in vivo teratoma model relative to control patient iPSCs. Furthermore, they found that treating mice bearing ACDC-iPSC teratomas with rapamycin reduced calcification (11).

As CD73-deficient mice do not sufficiently recapitulate the MAC observed in ACDC patients, we used the *Mgp*^-/-^ mice as an in vivo model of MAC to test the hypothesis that rapamycin could reduce vascular calcification in vivo. *Mgp*^+/+^ and *Mgp*^-/-^ mice were administered 5 mg/kg rapamycin or the same volume of vehicle (DMSO) three times per week starting at 10 days old until natural death (Figure 2A). We observed that rapamycin treatment extended the lifespan of *Mgp*^-/-^ mice (T_50_ 61 days) compared to the *Mgp*^-/-^ mice treated with the vehicle (T_50_ 35 days), which start to die as early as day 20 (Figure 2B). Both male (n=11) and female (n=8) mice were used; however, no sex differences were observed between *Mgp*^-/-^ mice that had received either vehicle or rapamycin treatment.

**Figure 2:**
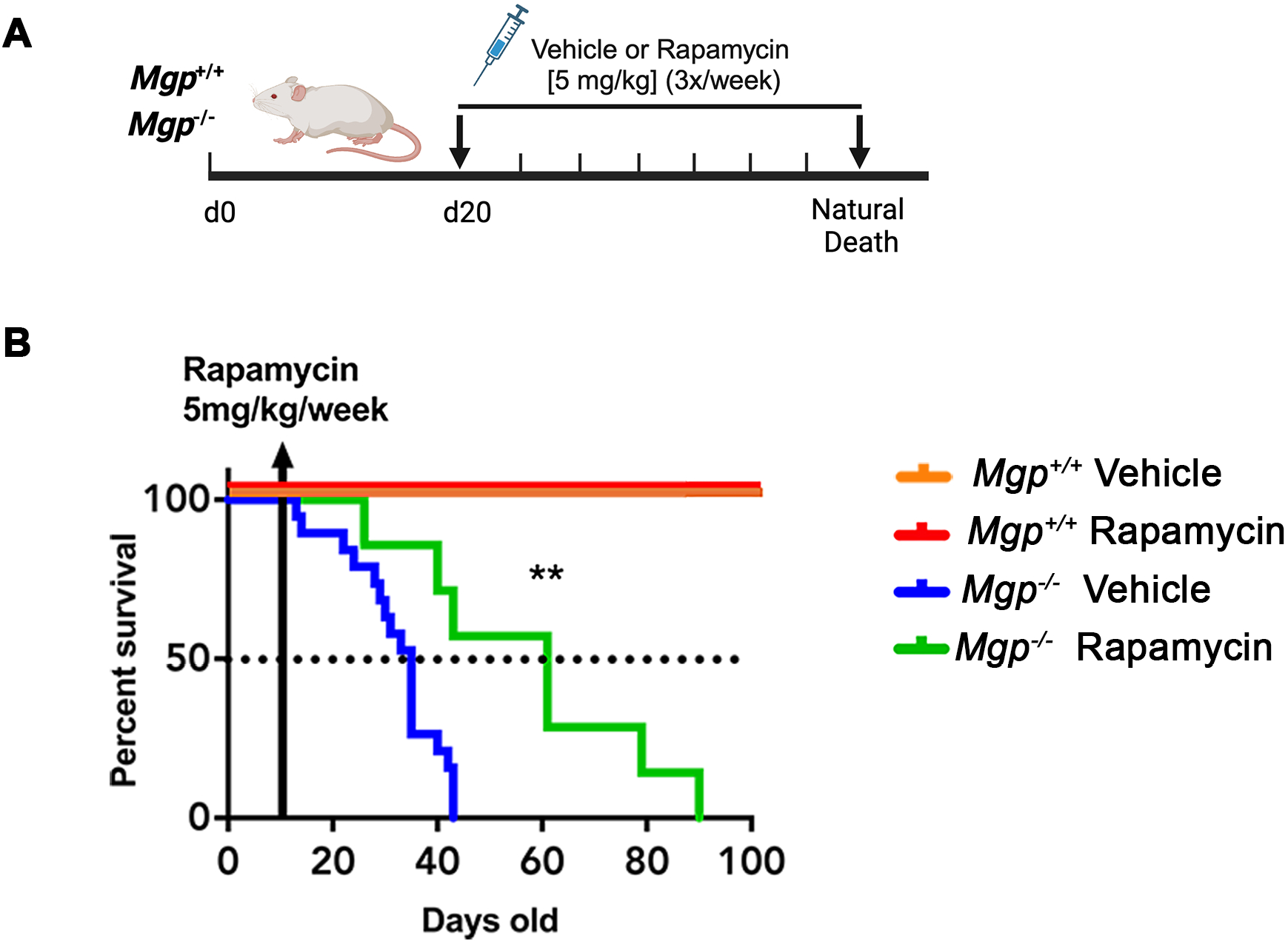
Impact of rapamycin treatment on survival in Matrix Gla Protein (*Mgp*)-deficient mice. **A**. Graphic illustration of vehicle (Dimethyl sulfoxide; DMSO) or rapamycin [5 mg/kg] treatment protocol. **B**. Kaplan-Meier analysis of overall survival in *Mgp*^*+/+*^ and *Mgp*^*-/-*^ mice treated with vehicle (DMSO) or rapamycin (5mg/kg/week) for 90 days until natural death. n=7 *Mgp*^*-/-*^ rapamycin, n=19 *Mgp*^*-/-*^ vehicle, n=8 *Mgp*^*+/+*^ rapamycin, n=5 *Mgp*^*+/+*^ vehicle. Mantel-Cox test and Gehan-Breslow-wilcoxon test were used for curve comparison.

We next investigated whether calcification content was altered in mice receiving rapamycin. Mice receiving vehicle, one (1x), or three doses (3x) of 5 mg/kg rapamycin per week, were sacrificed at day 25, and mineral content was assessed via microCT (Figure 3A). We selected the 25-day time point because we observed that *Mgp*^-/-^ mice start dying around that time (Figure 2B). Whole aorta microCT analysis found no differences in the total calcified deposits volume under either treatment regime; however, mice receiving rapamycin 3x per week exhibited a decreased mineral density compared to the control group (Figure 3B). Based on this data, we proceeded with 3x/week for the remaining studies. Von Kossa stain showed that the aortas of *Mgp*^*+/+*^ mice exhibit no mineralization, while both vehicle- and rapamycin-treated *Mgp*^-/-^ mice showed extensive calcification compared to *Mgp*^*+/+*^ mice. There were no discernible differences in the calcified area or the medial thickness of vehicle or rapamycin groups of *Mgp*^-/-^ (Figure 3C). RUNX2 is a key transcription factor promoting vascular cell osteogenic differentiation (6). Immunofluorescent staining revealed that while *Mgp*^-/-^ mice showed elevated levels of Runx2 compared to *Mgp*^*+/+*^ mice, rapamycin treatment did not reduce Runx2 protein levels in *Mgp*^-/-^ mice compared to *Mgp*^-/-^ mice treated with vehicle (Figure 3D). These findings suggest that while rapamycin doubles the lifespan of *Mgp*^-/-^ mice, it does not lead to decreased calcification volume, but does reduce the density of mineralized deposits in *Mgp*^-/-^ mice.

**Figure 3:**
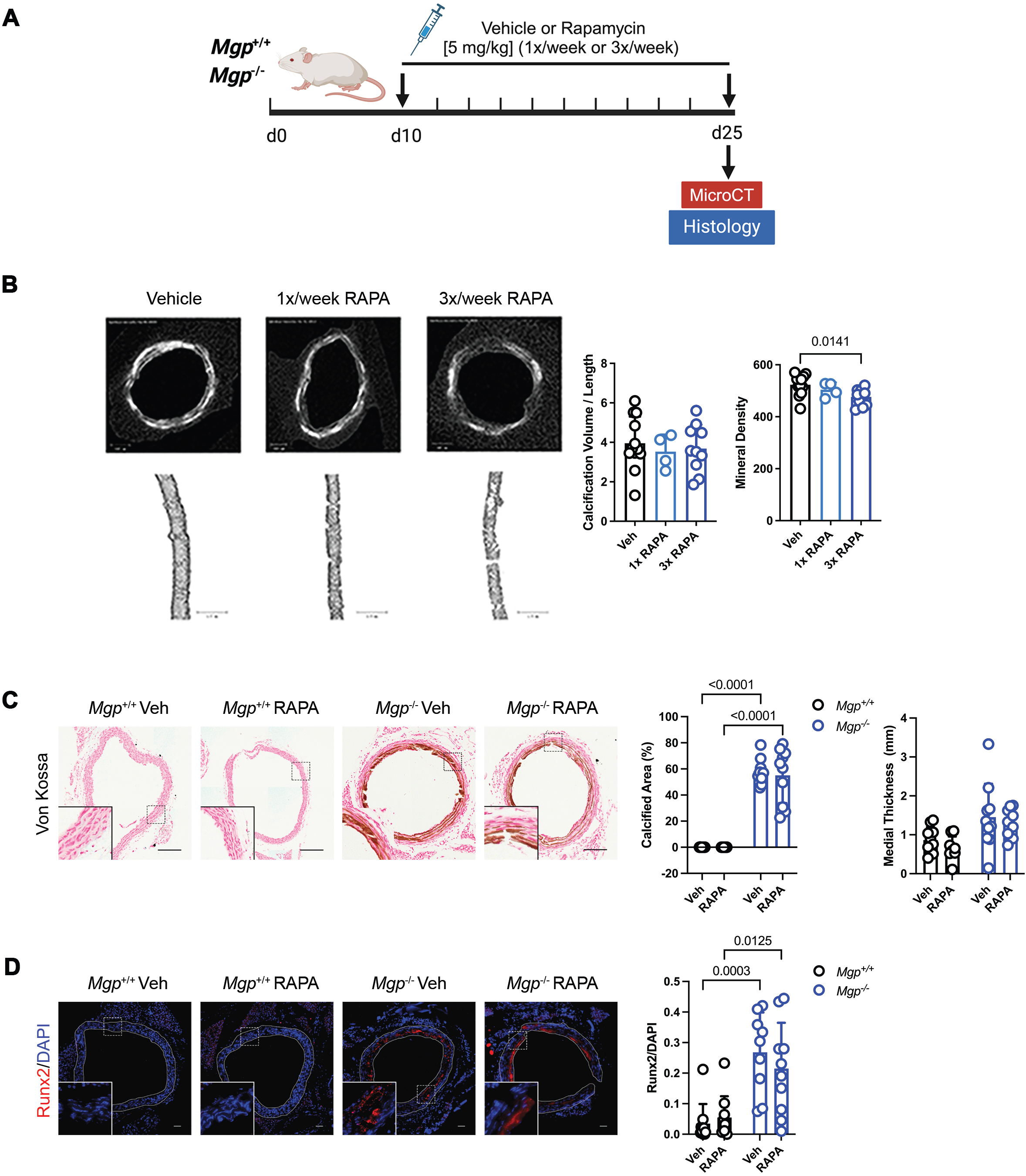
Effect of rapamycin treatment on medial calcification in *Mgp*-deficient mice. **A**. Graphic illustration of vehicle (DMSO) or 5 mg/kg rapamycin treatment for 25 days in *Mgp*^*+/+*^ and *Mgp*^*-/-*^ mice. **B**. Representative microCT measurement of axial and cross sections of *Mgp*^*-/-*^ mice aortas treated with vehicle or 5 mg/kg rapamycin, administered either once or three times a week for 25 days. Scale bar 0.2 mm. n=4 *Mgp*^*-/-*^ 1x rapamycin, n=10 *Mgp*^*-/-*^ 3x rapamycin, n=12 *Mgp*^*-/-*^ vehicle **C**. Representative Von Kossa staining images of *Mgp*^*+/+*^ and *Mgp*^*-/-*^ mice aortas treated with vehicle or rapamycin three times a week for 25 days. Scale bar 0.2 mm. n=10 *Mgp*^*-/-*^ rapamycin, n=10 *Mgp*^*-/-*^ vehicle, n=10 *Mgp*^*+/+*^ rapamycin, n=10 *Mgp*^*+/+*^ vehicle. **D**. Representative immunofluorescent images of Runx2 on the *Mgp*^*+/+*^ and *Mgp*^*-/-*^ mice aortas treated with vehicle or rapamycin three times a week for 25 days. Scale bar 50 μm. n=10 *Mgp*^*-/-*^ rapamycin, n=9 *Mgp*^*-/-*^ vehicle, n=10 *Mgp*^*+/+*^ rapamycin, n=10 *Mgp*^*+/+*^ vehicle. Data are shown as means ± SD. P values were calculated using Kruskal-Wallis test (Figure B) and Two-way ANOVA test (Figures C, D).

### Rapamycin alters SMC phenotype and ECM content in vivo

Rapamycin has a broad spectrum of effects on SMCs. Rapamycin inhibits SMC migration and proliferation by inducing the cyclin-dependent kinase inhibitors p27^kip^ and p21^cip^ to promote G1-S cell cycle arrest (35, 36). Rapamycin also induces SMC differentiation by inhibiting the mTOR-target S6K1 via AKT activation, promoting a contractile phenotype by enhancing the protein levels of SMC contractility markers smooth muscle α-actin (α-SMA) and myosin heavy chain 11 (Myh11) (37). Further, rapamycin was shown to protect against calcification of vascular cells in in vitro and ex vivo models (11, 23). While rapamycin did not reduce calcification volume in *Mgp*^*-/-*^ vessels we hypothesized that it could perhaps prevent aortic rupture by maintaining SMC contractile phenotype or stabilize the ECM.

Masson’s trichrome staining shows that *Mgp*^-/-^ vehicle vessels are highly remodeled, with acellular areas exhibiting observable increased collagen staining (Figure 4A). Picrosirius staining enabled the quantification of collagen content and revealed that thicker mature collagen in the red-orange colors and thinner immature collagen in green-yellow. In both the vehicle and the rapamycin treatment groups, the *Mgp*^*-/-*^ exhibited higher levels of both mature and immature collagen compared to the cognate *Mgp*^*+/+*^ mice. Within the *Mgp*^*-/-*^ genotype, rapamycin treatment reduced both types of collagen deposition compared to vehicle control (Figure 4B). This suggests that rapamycin helps restore or stabilize collagen homeostasis within the ECM, potentially preventing excessive fibrosis and maintaining tissue integrity (38), which may improve resilience against rupture. Verhoeff-Van Gieson (VVG) staining revealed that the aortas of *Mgp*^-/-^ mice exhibited less robust staining for elastin and less tortuous fibers in *Mgp*^-/-^ compared to *Mgp*^+/+^. While rapamycin did not alter elastic fiber tortuosity in *Mgp*^-/-^ mice compared to vehicle controls, the number of elastin breakages was higher in *Mgp*^*-/-*^ mice compared to *Mgp*^*+/+*^ in both groups and rapamycin treatment reduced the number of breakages in *Mgp*^*-/-*^ mice compared to vehicle (Figure 4C).

**Figure 4:**
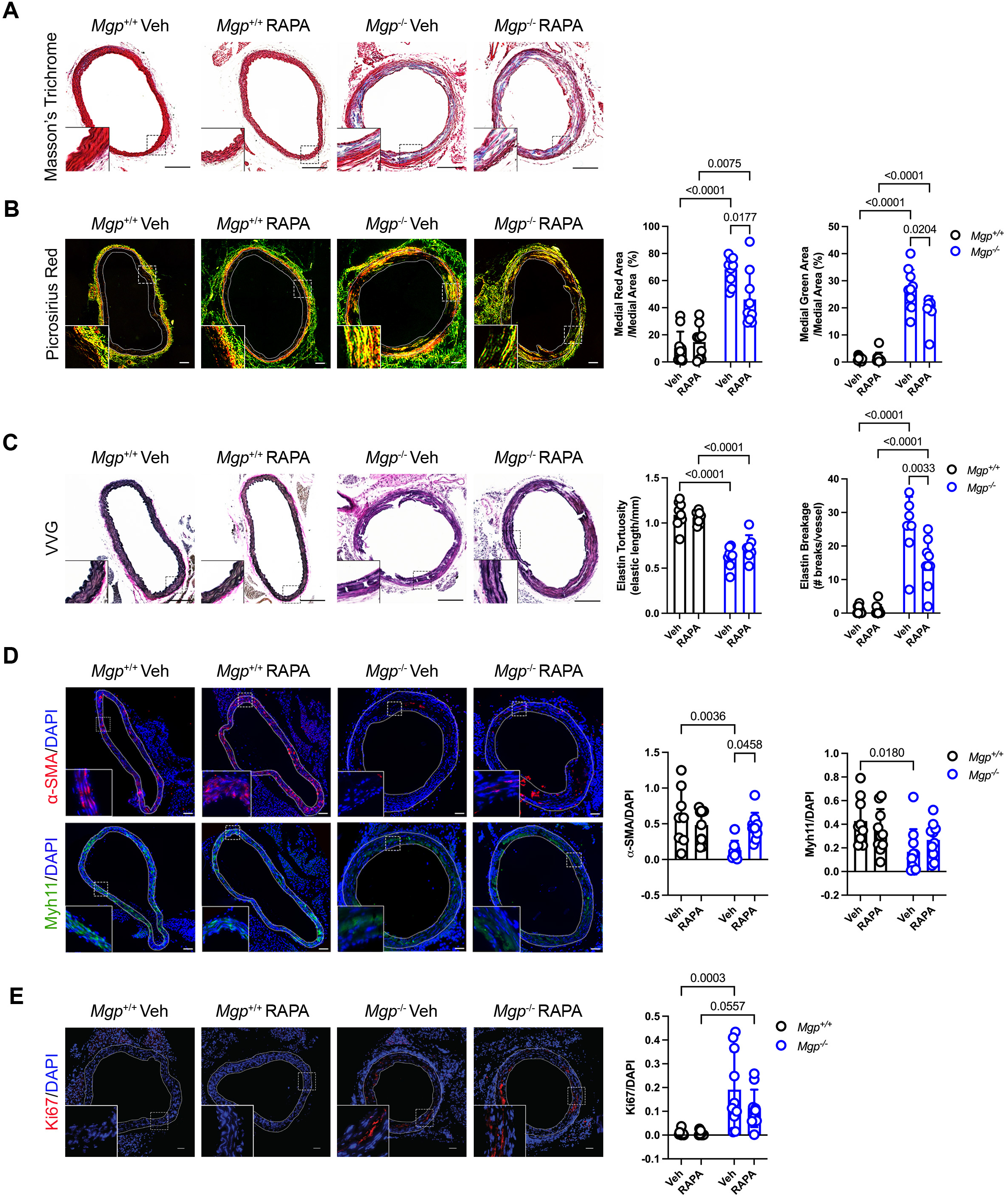
Histological structure of *Mgp*^*+/+*^ and *Mgp*^*-/-*^ mice aorta treated with vehicle or rapamycin. **A**. Representative Masson’s Trichrome staining images of *Mgp*^*+/+*^ and *Mgp*^*-/-*^ mice aortas treated with vehicle (Veh; DMSO) or rapamycin (RAPA; [5 mg/kg]) three times a week for 25 days. Collagen fiber structure in blue. Scale bar 0.2mm. **B**. Representative Picrosirius staining images of collagen in *Mgp*^*+/+*^ and *Mgp*^*-/-*^ mice treated with vehicle or rapamycin three times a week for 25 days. Scale bar: 50 μm. n=10 *Mgp*^*-/-*^ rapamycin, n=10 *Mgp*^*-/-*^ vehicle, n=10 *Mgp*^*+/+*^ rapamycin, n=10 *Mgp*^*+/+*^ vehicle. **C**. Representative Verhoeff-Van Gieson (VVG) staining images of *Mgp*^*+/+*^ and *Mgp*^*-/-*^ treated with vehicle or rapamycin three times a week for 25 days. Scale bar 0.2mm. n=7 *Mgp*^*-/-*^ rapamycin, n=8 *Mgp*^*-/-*^ vehicle, n=8 *Mgp*^*+/+*^ rapamycin, n=9 *Mgp*^*+/+*^ vehicle. **D**. Representative immunofluorescent images of smooth muscle actin (α-SMA) and myosin heavy chain gene (Myh11) SMA on *Mgp*^*+/+*^ and *Mgp*^*-/-*^ mice aortas treated with vehicle or rapamycin three times a week. Scale bar 50 μm. n=8 *Mgp*^*-/-*^ rapamycin, n=9 *Mgp*^*-/-*^ vehicle, n=10 *Mgp*^*+/+*^ rapamycin, n=10 *Mgp*^*+/+*^ vehicle. **E**. Representative immunofluorescent images of proliferation markers (ki67) on *Mgp*^*+/+*^ and *Mgp*^*-/-*^ mice aortas treated with vehicle or rapamycin three times a week for 25 days. Scale bar 50 μm. n=10 *Mgp*^*-/-*^ rapamycin, n=10 *Mgp*^*-/-*^ vehicle, n=10 *Mgp*^*+/+*^ rapamycin, n=10 *Mgp*^*+/+*^ vehicle. Data are shown as means ± SD. P value were calculated using Two-way ANOVA test.

Immunofluorescent staining was used to quantify SMC contractile proteins α-SMA and Myh11. In *Mgp*^*+/+*^ we observed no differences in α-SMA levels between vehicle and rapamycin treatment; however with vehicle treatment, *Mgp*^*+/+*^ mice exhibit higher α-SMA than *Mgp*^-/-^ mice. Rapamycin treatment rescued α-SMA levels in *Mgp*^-/-^ mice compared to vehicle alone, such that no differences were observed between *Mgp*^-/-^ mice treated with rapamycin and the *Mgp*^*+/+*^ vehicle or *Mgp*^*+/+*^ rapamycin groups. Similarly, *Mgp*^-/-^ mice treated with the vehicle exhibit reduced Myh11 levels compared to *Mgp*^+/+^ mice treated with the vehicle (Figure 4D). These results suggest rapamycin promotes the contractile phenotype of SMCs of *Mgp*^-/-^ mice. As the medial layer is remodeled in the *Mgp*^-/-^ mice, we also measured the mitotic marker Ki67 and found that at baseline, Ki67 levels were significantly elevated in *Mgp*^-/-^ aorta while nearly undetectable in *Mgp*^*+/+*^. However, rapamycin treatment did not alter Ki67 levels in either of the two genotypes (Figure 4E). Together, these data suggest that rapamycin promotes beneficial medial layer remodeling and a more contractile SMC phenotype in *Mgp*^-/-^ mice aorta.

### Rapamycin lowered LC3 levels in *Mgp*^*-/-*^ aorta, but there were no observable changes in autophagy flux

Autophagy is a ubiquitous process and contributes to bone development and the differentiation of osteoblasts and osteoclasts (39). One study found that autophagy reduces vascular calcification by limiting the release of pro-calcific matrix vesicles (40). Another found that in vitro autophagy stimulation with valproic acid protects against phosphate-induced calcification, while its inhibition increases extracellular vesicle release in SMCs (25). LC3 is a protein that functions at the initiation of autophagosome formation and serves as an autophagy marker. LC3 signal was significantly higher in *Mgp*^*+/+*^ aorta at baseline compared to *Mgp*^*-/-*^ mice. Within each genotype, rapamycin treatment significantly lowered LC3 puncta in *Mgp*^*+/+*^ compared to *Mgp*^*-/-*^ mice (Figure 5A).

**Figure 5:**
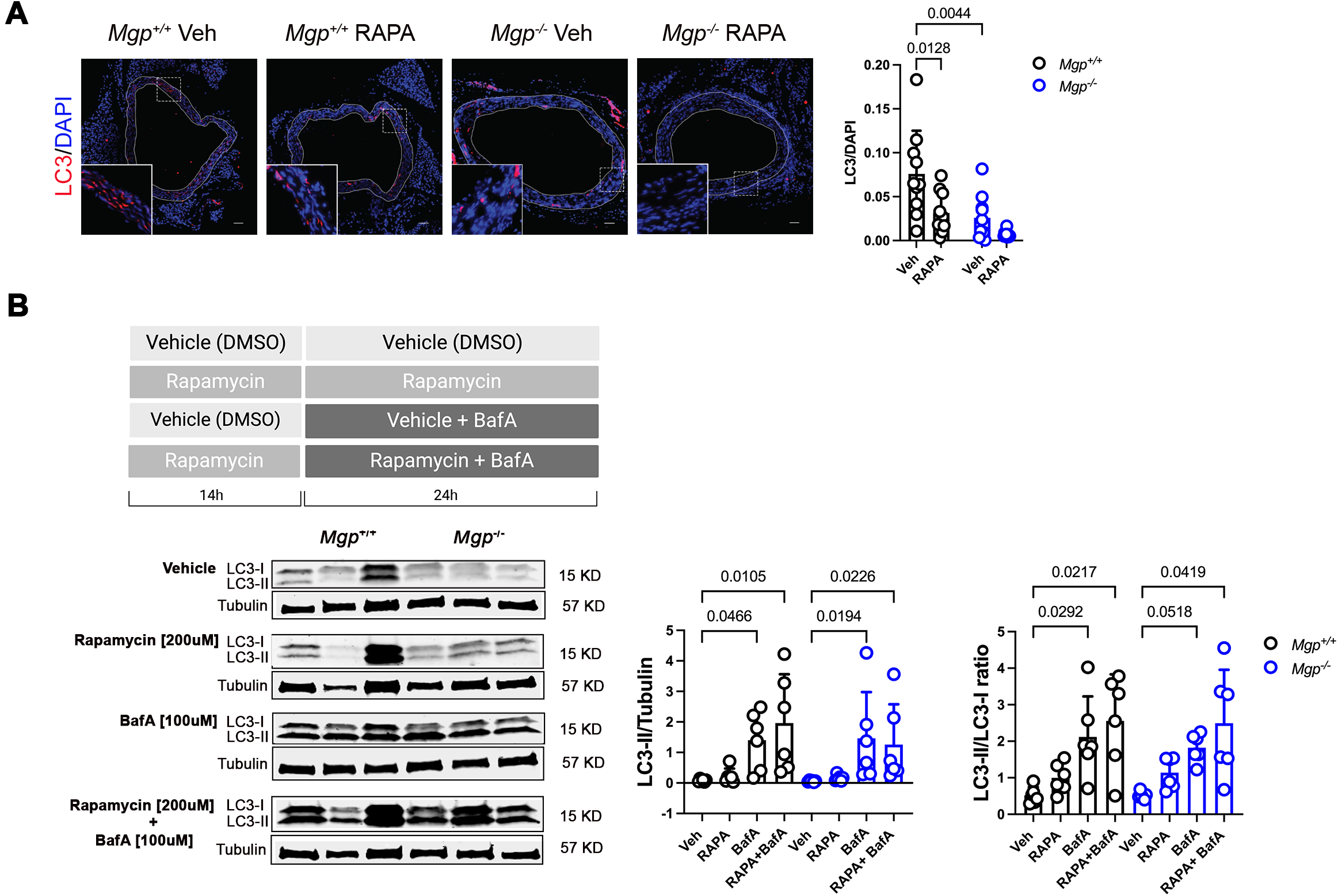
The effect of rapamycin treatment autophagy flux. **A**. Representative immunofluorescent images of LC3 on *Mgp*^*+/+*^ and *Mgp*^*-/-*^ mice aortas treated with vehicle (Veh; DMSO) or rapamycin (RAPA; [5 mg/kg]) three times a week for 25 days. LC3 decreased with rapamycin in *Mgp*^*-/-*^ mice compared to the *Mgp*^*+/+*^ mice. Scale bar 50 μm. n=9 *Mgp*^*-/-*^ rapamycin, n=10 *Mgp*^*-/-*^ vehicle, n=10 *Mgp*^*+/+*^ rapamycin, n=10 *Mgp*^*+/+*^ vehicle. **B**. Smooth muscle cells from *Mgp*^*+/+*^ and *Mgp*^*-/-*^ mice were treated with vehicle (DMSO) or rapamycin (200 µM) for 14 hours, followed by the addition of bafilomycin (BafA, 100 µM), an autophagy inhibitor, for an additional 24 hours. Levels of LC3 (LC3-I and LC3-II) were quantified by western blot. n=6 *Mgp*^*-/-*^ rapamycin, n=6 *Mgp*^*-/-*^ vehicle, n=6 *Mgp*^*+/+*^ rapamycin, n=6 *Mgp*^*+/+*^ vehicle. Data are shown as means ± SD. P values were calculated using Two-way ANOVA test (Figure A) and Kruskal-Wallis test (Figure B).

The quantification of LC3 puncta hints at diminished autophagy, but immunofluorescent staining of fixed tissues cannot assess autophagy flux because it provides only a single snapshot of the autophagic process, which does not reflect changes in the dynamic turnover of autophagosomes or the lysosomal clearance of LC3-positive autophagic vesicles. LC3-I is cleaved to LC3-II when it is incorporated into autophagosomes, but this process is dynamic, and LC3-II can also be degraded during autophagy. The comparison of LC3-II to LC3-I in the presence of an autophagy inhibitor like bafilomycin enables autophagy flux observation (41). We isolated SMCs from *Mgp*^*+/+*^ and *Mgp*^-/-^ mice to evaluate autophagy flux. SMCs were treated with vehicle (DMSO) or 200 nM rapamycin for 14h and then with vehicle or 100 uM or the late-phase autophagy inhibitor bafilomycin for 24h. Immunoblot analysis revealed that rapamycin alone did not increase LC3-II levels compared to vehicle controls in either the *Mgp*^*+/+*^ and *Mgp*^-/-^ SMCs. Bafilomycin increased LC3-II levels in both *Mgp*^*+/+*^ and *Mgp*^-/-^ SMCs compared to their cognate vehicle controls, however the addition of rapamycin did not further alter LC3-II levels compared to bafilomycin (Figure 5B, left graph). Comparing LC3-II to LC3-I showed identical trends (Figure 5B, right graph), suggesting that global LC3 levels and flux are similar in *Mgp*^*+/+*^ and *Mgp*^-/-^ SMCs.

### Genetic deletion of mTOR signaling in the SMCs does not replicate rapamycin treatment *Mgp*^-/-^ mice

Figure 2 shows that rapamycin treatment doubles the lifespan of *Mgp*^*-/-*^ mice compared to vehicle alone. However, our subsequent inquiries revealed that rapamycin did not decrease the volume of calcification as was seen in in vivo iPSC teratoma models (11), while it did promote less dense mineral formation, and beneficial SMC phenotype and EMC remodeling, albeit not to levels seen in the *Mgp*^*+/+*^ animals. mTOR can form two different protein complexes known as mTORC1, bound to Raptor, and mTORC2, bound to Rictor. The former is acutely sensitive to rapamycin administration, while the latter requires chronic rapamycin dosing for inhibition (21). To investigate whether rapamycin’s effects on the *Mgp*^-/-^ mouse are acting through either of these complexes, we generated *Mgp*^*-/-*^ mice with SMC-specific deletion of the mTOR complex proteins Raptor and Rictor (28-30). Fully differentiated SMCs expressing *Myh11* will constitutively express Cre recombinase to inactive *Raptor* and *Rictor* gene expression specifically in SMCs. *Mgp*^*-/-*^;*Myh11Cre-GFP;Raptor*^*f/f*^ mice (*Mgp*^*-/-*^;*Raptor*^SMC-/-^) were bred to examine the effects of a nonfunctional mTORC1 complex, while *Mgp*^*-/-*^;*Myh11Cre-GFP;Rictor*^*f/f*^ (*Mgp*^*-/-*^;*Rictor*^*SMC-/-*^) mice knockout was bred to examine a nonfunctional mTORC2 complex.

Similar to *Mgp*^*-/-*^ mice treated with vehicle in Figure 2, Figure 6 shows that only 50% of the *Mgp*^*-/-*^ ;*Rictor*^*SMC-/-*^ and *Mgp*^*-/-*^;*Rictor*^*SMC+/+*^ are alive at 40 days, indicating that rapamycin acting via mTORC2 does not recapitulate the effects of rapamycin on increasing *Mgp*^*-/-*^ mice lifespan. The 50% survival rate of *Mgp*^*-/-*^;*Raptor*^*SMC+/+*^ mice was similar to *Mgp*^-/-^ treated with vehicle, *Mgp*^*-/-*^ ;*Rictor*^*SMC-/-*^ and *Mgp*^*-/-*^;*Rictor*^*SMC+/+*^, however the 50% survival rate of *Mgp*^*-/-*^;*Raptor*^*SMC-/-*^ mice dropped to 22 days. As the Cre strain being used is a non-inducible constitutive expression system, this data from *Mgp*^*-/-*^;*Raptor*^*SMC-/-*^ mice suggest that mTORC1 is critical to the growth and maturation of the vasculature post birth. Together, these in vivo data suggest that the life-extending effects of rapamycin in the *Mgp*^*-/-*^ mouse are not acting through the mTORC2 complex. Additional experiments, such as a tamoxifen inducible Cre system, would need to be used to fully address the contribution of mTORC1 to mirror effects seen by rapamycin injection.

**Figure 6:**
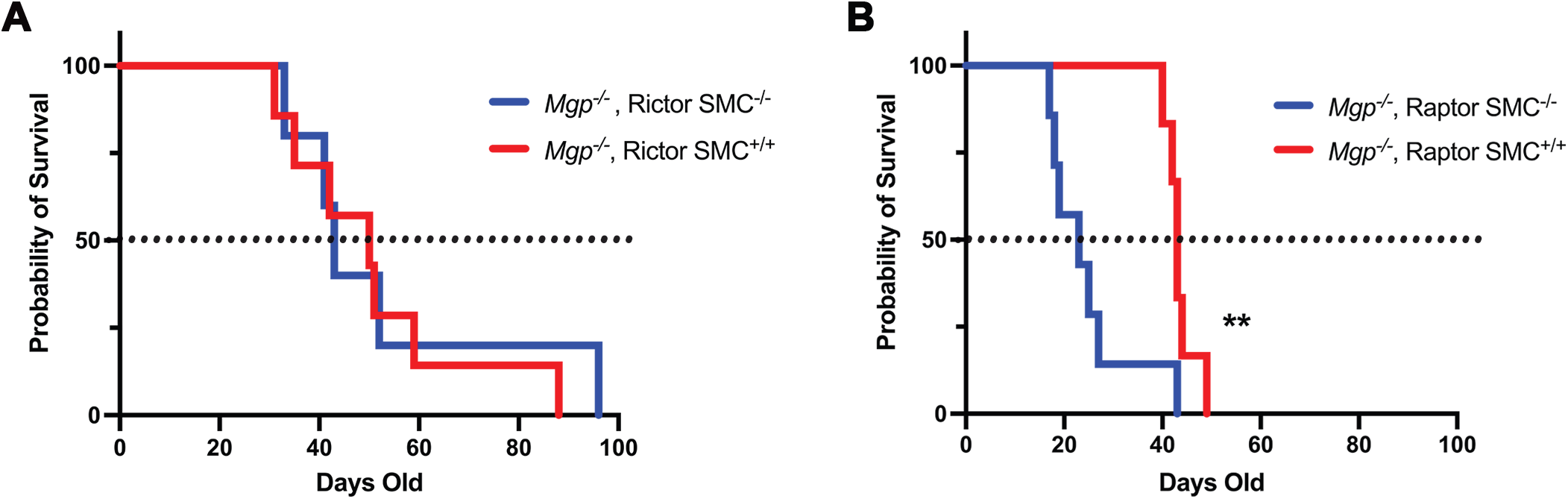
Survival probability of Matrix Gla Protein (MGP)-deficient mice with Rictor and Raptor knockouts in vascular smooth muscle cells (SMC). **A**. *Mgp*^*-/-*^ using Cre recombinase that targets the gene encoding Rictor in SMC, a protein involved in mTORC2 signaling pathways. *Mgp*^*-/-*^;*Rictor*^*SMC-/-*^ (n=5) and *Mgp*^*-/-*^;*Rictor*^*SMC+/+*^ (n=7). **B**. *Mgp*^*-/-*^ using Cre recombinase that target the gene encoding Raptor in SMC a protein involved in mTORC1 signaling pathways. *Mgp*^*-/-*^;*Raptor*^*SMC+/+*^ (n=6), *Mgp*^*-/-*^;*Raptor*^*SMC-/-*^ (n=7). Data are shown as means ± SD. Mantel-Cox test and Gehan-Breslow-wilcoxon test were used for curve comparison.

## DISCUSSION

In this study, we demonstrated that rapamycin treatment increased the lifespan in *Mgp*^-/-^ mice, but rapamycin does not reduce MAC per se; while mineral density is lower with rapamycin treatment the mineral volume remains unchanged. In mice, rapamycin has been shown to increase maximum lifespan and delay the onset of cancer as hyperactivity of the mTOR pathway leads to the expression and activation of many oncogenes like PI3K, Akt, and eIF4E. Hence, rapamycin can slow the proliferation of tumor cells by causing cell arrest, promoting apoptosis, and blocking angiogenesis in tumor growths (42). Our data show that the increase in lifespan observed in *Mgp*^*-/-*^ mice following rapamycin treatment is associated with improvements in SMC contractile protein expression and a normalization of collagen content in the vessel wall, but not with enhanced autophagy flux.

Collagen, the most abundant protein in the ECM, plays a crucial role in maintaining the structural integrity of the blood vessels. Collagen dysregulation or degradation leads to increased vessel distensibility and the risk of rupture (43, 44). Aging mice homozygous for a deletion in the gene coding for collagen alpha 1(*Col1a1*) are at a significantly increased risk of aortic dissection and rupture (45). Our results align with a study which showed elevated collagen deposition in *Mgp*^*-/-*^ mice aorta (18). In 25-day-old *Mgp*^*-/-*^ mice, rapamycin treatment reduced collagen content in the medial layer compared to vehicle-treated mice, bringing it to levels observed in *Mgp*^*+/+*^. Total adventitial collagen content in the mice aorta increases with age (46), suggesting the differential distribution we observed was due to the young age of the mice in our study. The decrease in collagen content following rapamycin treatment and the increase in SMC contractile protein SMA may contribute to the prolonged lifespan of *Mgp*^*-/-*^ mice by improving both the structural stability and functional resilience of the vascular walls.

While rapamycin is known to inhibit SMC proliferation and is used in drug-eluting stents, there is limited research on the effects of rapamycin on vascular morphology in the presence of calcification (36, 47). It was previously shown that treatment with rapamycin limits the progression of abdominal and thoracic aortic aneurysms in mice and preserves elastic lamina integrity (48, 49). In a hyperphosphatemic (high-phosphate, high-adenine diet) rat model, rapamycin treatment inhibited aortic calcium deposition and reduced the osteogenic markers *Msx2* and *Osx* while increasing the *Opn* and *Acta2* gene expression via Klotho upregulation (50). Opposing this diet-induced model of calcification, we did not observe a rapamycin-induced reduction of the osteogenic marker Runx2 in *Mgp*-deficient mice. Our approach used a genetic model of MAC to investigate whether rapamycin could inhibit calcification in vivo. *Mgp*-deficient mice develop normally to birth without any overt abnormalities. At approximately one week old *Mgp*^*-/-*^ mice start exhibiting calcification of the vessels, suggesting that other gene products might compensate to prevent calcification during development and immediately post-birth (18).

These observations highlight the differences between genetically induced MAC and high phosphate feeding models. Diet-induced calcification must overcome the effect of endogenous Mgp, which is a potent inhibitor of calcification, while the genetic removal of *Mgp* results in calcification from the circulating calcium and phosphate and activation of BMP signaling (14, 17).

Rapamycin has also been shown to preserve the differentiated phenotype of vascular SMCs through IRS-1/P3K/AKT2 pathway (37). Our study did observe that rapamycin enhanced the expression of a SMC contractile proteins in *Mgp*^-/-^ mice. *Sma* expression typically occurs earlier than *Myh11* expression during SMC development (51). In contrast, *Myh11* expression occurs later, during the maturation of SMCs. One study examined SMC contractile gene expression in response to rapamycin in vitro (37), while our study looked at the SMC contractile markers in vivo. Further, the presence of calcification activates mechanosensing pathways that once activated may prevent full re-expression of SMC contractile markers (52). However, further investigation is needed to elucidate how these mechanisms are operating in the context of the *Mgp*^-/-^ mouse.

MAC is characterized by mineral deposits in the medial vessel layer and frequently occurs in patients with PAD. Vascular calcification is an active biological process with many driving forces, including osteogenic dedifferentiation of SMCs, inflammatory signals, hyperphosphatemia, and extracellular matrix remodeling (53). In this study, rapamycin prolonged the lifespan of mice who developed significant MAC shortly after birth. Interestingly, even though rapamycin has been shown to stimulate the cellular process of autophagy (54)— which can recycle components of the ECM, including mineral deposits (55)—calcification volume was not reduced in rapamycin treated *Mgp-/-*mice. However, microCT data showed that the mineral density was significantly reduced in *Mgp*^*-/-*^ mice treated with rapamycin. The decrease in mineral density without a change in calcification volume could suggest a shift from compressed, organized macrocalcifications to more dispersed calcium-rich extracellular vesicles and amorphous calcium phosphate deposits (7, 56). Amorphous calcium phosphate, primarily composed of calcium and phosphate ions, represents the early stages of calcification (57). It subsequently acts as a precursor for the nucleation and growth of hydroxyapatite crystals within the ECM. Stabilization of hydroxyapatite crystals further contributes to the transition into a bone-like phenotype. One reason that a lower mineral density did not affect total calcification volume could be a difference in the pattern of mineral deposition. For example, the mineral may form in a less compact, more porous manner, leading to a lower overall density despite occupying the same spatial volume. Alternatively, the reduction in density could result from the breakdown or remodeling of pre-existing mineral deposits. This suggests that changes in mineral structure, if explored at a further timepoint may impact calcification volume. Alternatively, as Mgp also functions to inhibit BMP2 signaling, rapamycin may not interfere with that process, such that changes in dosage or duration of rapamycin treatment would still not alter calcification content in the vessel wall. Another explanation is that the activated autophagy pathways cannot overcome the rapid deposition of hydroxyapatite that occurs in *Mgp*^-/-^ mice. Frauscher et al. demonstrated that induction of autophagy by rapamycin treatment decreased calcification in the vasculature of uremic DBA/2 mice (58). This argument, along with our prior data from iPSC-based disease modeling (11), led us to examine autophagy flux in SMCs. However, we did not observe a difference in autophagy flux between *Mgp*^*+/+*^ and *Mgp*^-/-^. It is important to note that the *Mgp*^-/-^ is on a C57BL6J background, perhaps explaining the differences observed in these in vivo models.

Rapamycin is an inhibitor of mTOR, but mTOR pathways are complex. mTOR forms two distinct protein complexes known as mTORC1 and mTORC2. mTORC1, activated by amino acids and allosterically inhibited by rapamycin, requires the protein named Raptor (59). mTORC1 is acutely sensitive to rapamycin, while only after prolonged rapamycin treatment (>6 hours) does mTORC2 (requiring Rictor) acquire a sensitivity to rapamycin (22, 60). In the *Mgp*^*-/-*^;*Raptor*^*SMC-/-*^ mice, induction of Cre to knockout *Raptor* was more lethal than the *Mgp*^*-/-*^;*Raptor*^*SMC+/+*^ or *Mgp-/-*mice. This suggests that *Raptor* is essential for the growth and maintenance of SMCs in the vasculature. This essential role in growth and development is perhaps unsurprising, considering that global *Raptor* knockout mice die in utero (61). Moreover, there was no benefit to lifespan in the *Mgp*^-/-^:Rictor^SMC-/-^ mice. The survival curves for the *Mgp*^*-/-*^:*Rictor*^*SMC+/+*^ and *Mgp*^*-/-*^;*Rictor*^*SMC-/-*^ mice, targeting mTORC2 signaling, do not diverge at any point and are statistically insignificant and also mirror that seen in the *Mgp*^-/-^ strain.

Arterial stiffness is strongly linked to heart failure (62), and these non-atherosclerotic vascular pathologies are much more common in some populations, such as African-ancestry individuals who are known to have a higher risk of cardiovascular disease events and mortality compared to Caucasians (63-67). This excess cardiovascular disease risk is largely due to hypertensive and peripheral vascular disease versus atherosclerotic disease (63-65, 68-70). As such, it is critical to define drivers of MAC such as MGP, which in some clinical studies in humans has been associated with heart failure and MAC (71). Impairment in MGP activation, which would phenocopy genetic deletion and lead to arterial stiffening and peripheral artery disease, is linked to a higher risk of heart failure due to chronically elevated afterload leading to left ventricle remodeling (72, 73). As the mice that lack *Mgp* are prone to aortic rupture or heart failure, we speculate that rapamycin’s effect on the mice’s lifespan may be due to rapamycin acting on the cardiac tissues.

Our study suggests that rapamycin’s life-extending benefit in *Mgp*^-/-^ mice may act through improvement in vascular function, which could prevent aortic rupture or through another organ system, possibly the heart. However, further studies are needed to examine the impact of rapamycin on cardiac function in *Mgp*^-/-^ mice. Rapamycin has been shown to increase cardiomyocyte autophagy (74, 75), limit cardiomyocyte death, and attenuate cardiomyocyte hypertrophy and cardiac remodeling by enhancing mTORC2 signaling while simultaneously inhibiting mTORC1 signaling (76). Rapamycin had a protective effect on cardiomyocytes isolated from the old mice compared to old mice not treated with rapamycin, and improved ejection fraction, reduced cardiac hypertrophy and inflammation, and downregulated ANP-induced stress response in vivo (77, 78). In mice with decompensated cardiac hypertrophy, rapamycin improved cardiac function, such as left ventricular end-systolic dimensions, fractional shortening, and ejection fraction in mice compared to the control group (79). Three months of rapamycin treatment in aged mice increased lifespan, decreased heart rate and heart weight (80). However, a calcified phenotype was not noted in any of these animal studies.

## CONCLUSION

This study provides novel insights into the effects of rapamycin on medial arterial calcification (MAC) using the *Mgp*-deficient murine model. While rapamycin did not reduce the volume of calcification in the vasculature, it significantly prolonged the lifespan of *Mgp*^*-/-*^ mice, possibly through stabilizing vascular structure. This was evidenced by reduced collagen content and enhanced SMC contractile protein expression, which collectively suggest improved resilience against vascular rupture. Interestingly, while the volume of calcification was not changed, rapamycin treatment reduced mineral density. Further, our study underscores the importance of distinguishing between mTORC1- and mTORC2-mediated pathways. Rapamycin’s life-extending effects appeared independent of autophagy flux or mTORC2 signaling, while the critical role of mTORC1 in vascular development and maintenance was highlighted, as *Mgp*^*-/-*^;*Raptor*^*SMC-/-*^ died earlier than *Mgp*^*-/-*^ mice. These findings illustrate rapamycin’s role in modifying calcification characteristics and enhancing vascular integrity without altering overall calcification volume. Future investigations should focus on elucidating rapamycin’s systemic effects, such as its impact on cardiac function and/or the immune system, as well as understanding how it leads to less-dense mineral deposits, to clarify its protective mechanisms in the context of MAC and cardiovascular health.

## ABBREVIATIONS

AMP: Adenosine monophosphate
ATP: Adenosine triphosphate
TNAP: Alkaline phosphate
ACDC: Arterial calcification due to deficiency of CD73
NT5E: Ecto-5’-nucleotidase
iPSCs: Induced pluripotent stem cells
Mgp: Matrix GLA protein
MAC: Medial arterial calcification
mTORC1: mTORC complex 1
mTORC2: mTORC complex 2
PAD: Peripheral artery disease
α-SMA: Smooth muscle α-actin
SMCs: Smooth muscle cells
Myh11: Myosin heavy chain 11
Runx2: Runt-related transcription factor 2

## ACKNOWLEDGEMENTS

We want to thank the histology core and the Center for Biologic Imaging at the University of Pittsburgh for their help and support with microscopy imaging. Some figures were created with Biorender.com.

## FIGURE LEGENDS

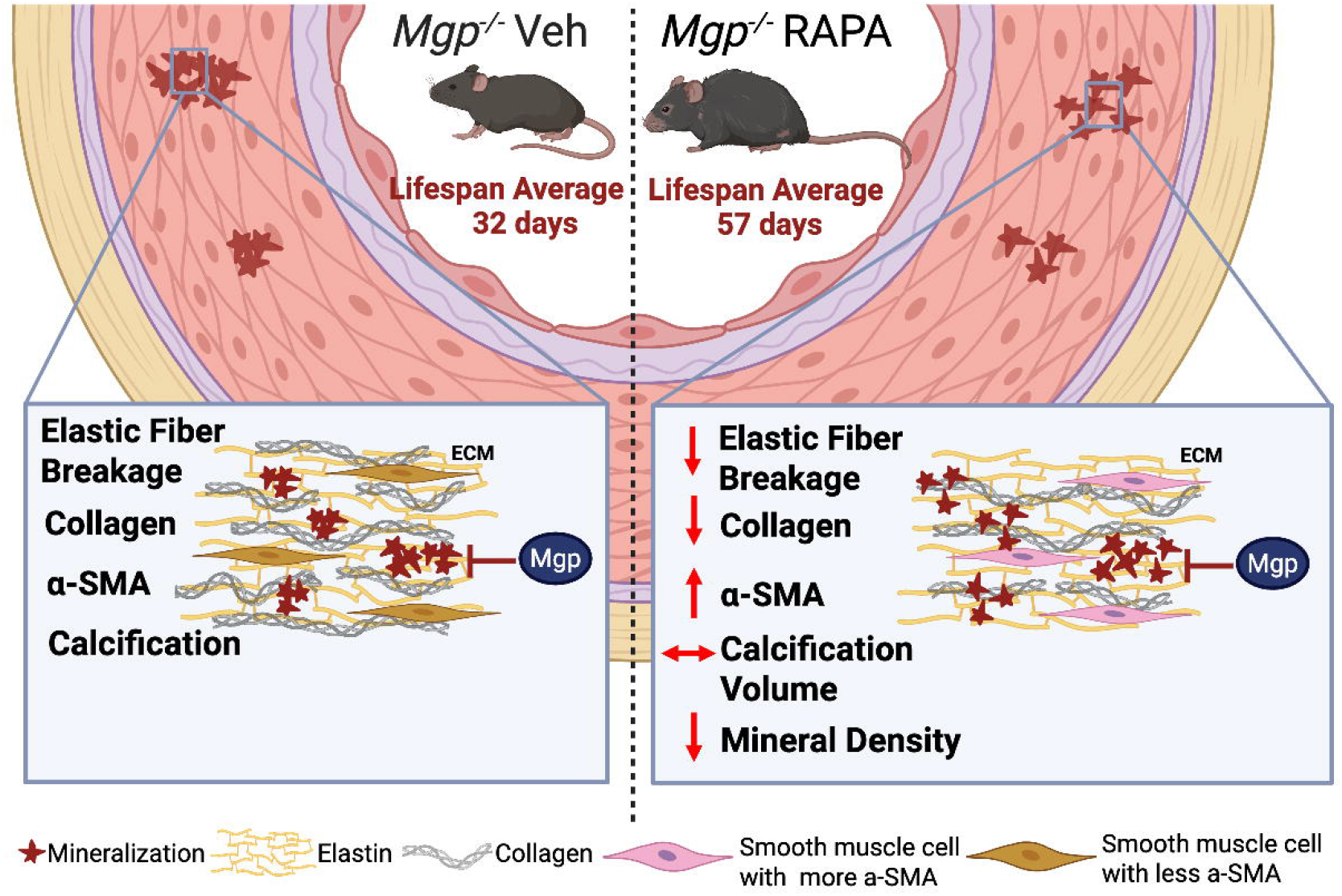

